# Upregulation of *Trem2* expression occurs exclusively on microglial contact with plaques

**DOI:** 10.1101/2022.01.26.477873

**Authors:** Jack Wood, Eugenia Wong, Ridwaan Joghee, Aya Balbaa, Karina S. Vitanova, Alison Vanshoiack, Stefan-Laural J. Phelan, Francesca Launchbury, Sneha Desai, Takshashila Tripathi, Jörg Hanrieder, Damian M. Cummings, John Hardy, Frances A. Edwards

**Affiliations:** Department of Neuroscience, Physiology & Pharmacology, University College London, Gower Street, London WC1E 6BT, United Kingdom; Nanostring Technologies, 530 Fairview Ave N, Seattle, WA 98109, United States; Department of Neurodegenerative Disease, University College London Queen Square Institute of Neurology, Queen Square, London WC1N 3BG, United Kingdom; Dementia Research Institute, University College London, Gower Street, London WC1E 6BT, United Kingdom; Department of Psychiatry and Neurochemistry, Institute of Neuroscience and Physiology, Sahlgrenska Academy at the University of Gothenburg, Mölndal, Sweden; Institute of Healthy Ageing, University College London, Gower Street, London WC1E 6BT, United Kingdom

**Keywords:** Microglia, NLF mouse, *App^NLF/NLF^*, *Trem2*, *Trem2^R47H^*, Spatial-Transcriptomics, PIGs, Alzheimer’s Disease, astrocytes, knock-in mouse

## Abstract

Using spatial cell-type-enriched transcriptomics, we compare plaque-induced gene (PIG) expression in microglia touching plaques, neighboring plaques, and far from plaques in 18-month-old *APP^NLF/NLF^* knock-in mice with and without the Alzheimer’s disease risk mutation *Trem2^R47H/R47H^*. We report that, in *App^NLF/NLF^* mice, expression of 35/55 PIGs, is exclusively upregulated in microglia that are touching plaques. In 7 PIGs including *Trem2* this upregulation is prevented by the *Trem2^R47H/R47H^* mutation. Unlike in young mice, knockin of the *Trem2^R47H/R47H^* mutation does not significantly decrease the *Trem2* expression but decreases protein levels by 20% in the absence of plaques. On plaques, despite the mutation preventing increased gene expression, TREM2 protein levels increased by 1.6-fold (compared to 3-fold with *Trem2^WT/WT^*) and microglial density increased 20-fold compared to 30-fold. Hence microglia must touch plaques before *Trem2* gene expression is increased but small changes in protein expression can increase microglia density without a change in gene expression.

## Introduction

The importance of the microglial membrane receptor TREM2 in Alzheimer’s disease (AD) is now well known. The initial evidence came from genome-wide association studies (GWAS), which convincingly demonstrated that *TREM2* has several variants that increase the risk of reaching the stage of AD diagnosis. *TREM2^R47H^*, the most common variant, has an effect size similar to that of the *APOE4* allele in people of white European descent (Gratuze et al., 2018; Guerreiro et al., 2013; Jonsson et al., 2013). Considerable work has been undertaken using mouse models carrying familial AD mutations, *Trem2* knock-out mice, and primary microglial cultures (Liu et al., 2020) (reviewed in Gratuze *et al*., 2018; Kulkarni et al., 2021), revealing that TREM2 pushes microglia towards an anti-inflammatory phagocytic phenotype and has been reported to be instrumental in the increased density of microglia around plaques. Moreover, microarray analysis of transgenic mice, carrying familial AD mutations, revealed *Trem2* and an array of co-expressed genes to be upregulated in the hippocampus and cortex (Matarin et al., 2015), a finding repeatedly confirmed with RNAseq, both in whole tissue (Salih et al., 2019) and in single-cell studies. Such single-cell RNAseq studies have confirmed patterns of gene expression defining “disease-associated microglia (DAM)” genes (Deczkowska et al., 2018; Keren-Shaul et al., 2017), in relation to amyloidβ (Aβ) and Tau pathology (Lee et al., 2021b). A recent study using spatial transcriptomics in relation to plaque density defined a further overlapping group of genes referred to as plaque-induced genes (PIGs, Chen et al., 2020).

Most of the mouse studies published have focussed on transgenic mice that overexpress familial AD mutations in the amyloid precursor protein (*APP*) and/or the presenilin *(PSEN)* genes. Moreover, when knock-in mice are used, the most popular model has been the *App*^NLGF/NLGF^ (NLGF) mouse (Saito et al., 2014) which harbours three familial AD mutations in *APP* and includes a humanised Aβ sequence. Similarly to transgenic models, NLGF mice deposit plaques rapidly and early, such that the most rapid increase in plaque load occurs between 2 and 4 months of age, reaching almost maximal density by 9-months (Benitez et al., 2021). Sporadic AD develops slowly from midlife into old age, and so the reaction of microglia to this rapid rise of plaques in young animals may be very different and less relevant than their response to slow deposition starting later in life. Consequently, we would suggest that the *App*^NLF/NLF^ (NLF) mouse (Saito *et al*., 2014) is a better model of sporadic AD than the NLGF mouse as in NLF mice, plaques begin to develop slowly in midlife and increase through to at least 24 months of age. Although using these very old mice is inconvenient and increases the cost of studies, the increased relevance of combining slow development and, importantly, the element of old age, outweighs these disadvantages. However, until recently, analysing changes in gene expression was very difficult in NLF mice because, even at 24 months of age, genes such as *Trem2* show only modest changes, despite evident synaptic differences and increased microglial density (Benitez *et al*., 2021). It is this combination of high-cost and low reward that has made NLFs a less popular model, despite the similarity to the progression of human sporadic AD. However, with the introduction of spatial cell-type-enriched transcriptomics, more subtle and direct analysis of plaque-induced microglial gene expression changes becomes possible, without the diluting effects of bulk analysis.

Using spatial transcriptomics in this more relevant mouse model, we initially compared microglial-enriched expression of ‘plaque-induced genes’ (PIGs). PIGs were defined by Chen et al. in NLGF mice as genes that upregulate in response to plaque density in different brain regions and across multiple cell types (Chen *et al*., 2020). We confirm in NLF mice that more than half of these genes were more highly expressed in microglia directly in contact with plaques than in those distant from plaques or in the brains of mice without plaques. Importantly, this only occurred in microglia directly in contact with plaques and not in immediately neighbouring regions.

We particularly study *Trem2* as one of the genes upregulated specifically on plaques in these mice. We compared these effects to NLF mice in which *Trem2^R47H/R47H^* is knocked in (NLFTrem2^R47H^ mice) and validated our findings at the protein level using immunohistochemistry. The lack of *Trem2* upregulation in the NLFTrem2^R47H^ mice could then be used to assess the dependence of increased microglial density at plaques on *Trem2* expression. Surprisingly, in NLFTrem2^R47H^ mice, the density of microglia increased considerably which may be due to a modest increase in TREM2 protein expression, in the absence of increased gene expression.

## Results

In NLF mice, plaques are first detected at 9 to 10 months of age. Deposition then slowly increases through to 24 months of age (Benitez *et al*., 2021). In the present study we investigate the spatial distribution of gene expression in 18-month-old NLF and NLFTrem2^R47H^ mice.

For spatial transcriptomics analysis, we labelled plaque pathology and associated glial cells using immunohistochemistry towards TMEM119 (microglia), GFAP (astrocytes) and Aβ40/Aβ42 (plaques) (Figure 1A). We then selected large regions of interest (ROIs) that were densely filled with plaques and nearby ROIs without plaques (Figure 1Bi and 1Bii). Most plaques in the hippocampus of NLF mice are found in and around the *stratum lacunosum moleculare* (SLM) of CA1-3 and the *stratum moleculare* of the dentate gyrus (Figure 1Bii). It is thus possible, in each brain section, to define a region in which all pixels are within ~30 μm of a plaque and other regions in which no plaques are detected. We then defined cell type-enriched areas of interest (AOIs) for microglia or astrocytes within each ROI. For the present study, we focussed on microglia and determined genome-wide RNA signatures from the following three AOIs within each ROI in NLF and NLFTrem2^R47H^ mice: 1. *<u>On plaque</u>*, all the microglia in the plaque ROIs that colocalised with Aβ (Figure 1Ci); 2. *<u>Periplaque</u>*, all the microglia in the plaque ROI that did not colocalise with Aβ (Figure 1Cii); 3. *<u>Away</u>* from plaque, all the microglia in the ROIs without plaques. In addition, ROIs in equivalent positions were studied in wild type (WT) and Trem2^R47H^ mice, so that the same regions could be compared without plaques. Before collecting RNA signatures from the microglial AOIs, signatures were collected from the regions labelled with GFAP in order to decrease contamination from overlapping astrocytes. However, it is important to note that the results represent enrichment for microglia rather than selective analysis of microglial genes. Indeed, some highly expressed genes from astrocytes and neurones are also detected in the microglial gene set. Nevertheless, a substantial degree of separation between cell types is achieved, as can be seen when comparing the expression of genes previously defined as microglia-enriched (Ximerakis et al., 2019)(Figure S1A) and in principal components analysis (Figure S1B).

**Figure 1.**
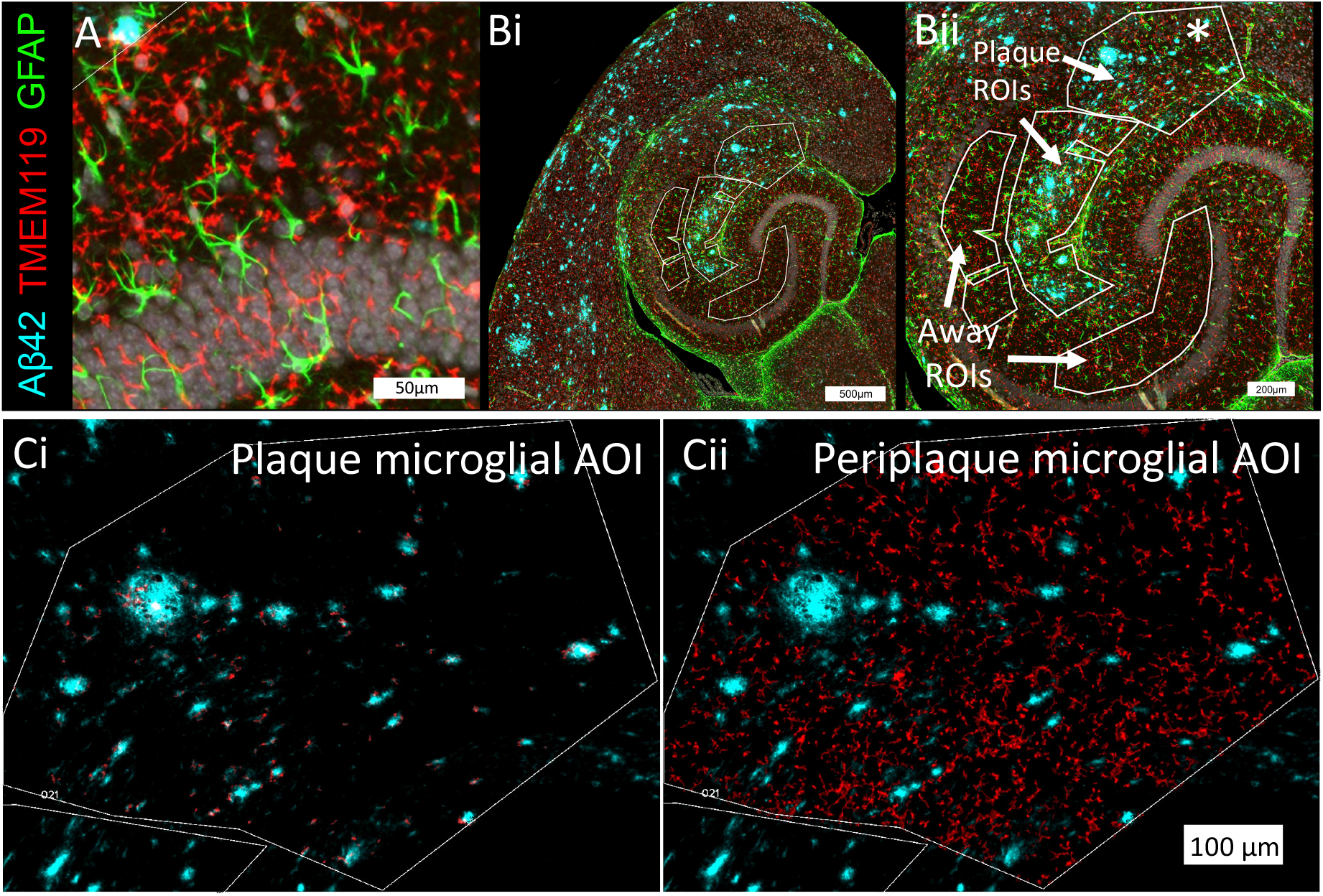
Selection of Areas of Interest (AOIs) Sections of the hippocampus and neighbouring cortex (8 μm thick) were mounted and labelled with antibodies for plaques (cyan, Aβ40/Aβ42), microglia (red, TMEM119), astrocytes (green, GFAP) and nuclei to reveal the cell bodies of the hippocampus (white, DAPI). **(A)**. Example image: microglia (red) and astrocytes (green) could be readily identified. A small plaque is visible in the top left corner (cyan) and a section of the dentate gyrus granule cell body layer is visible in white. **(B)** Regions of interest were established that either contained a heavy plaque load (Plaque ROI) or lacked plaques (Away ROI). Bii shows a zoomed in image of the hippocampal region of the full section shown in Bi. * indicates the ROI detailed in C. **(C)** To maximise the separation of microglia, the samples from astrocytes (GFAP labelled areas) were first removed before collecting from the microglia regions. The plaque ROIs were then separated into sub areas of interest (AOIs) whereby the Plaque microglial AOI was defined as TMEM119 positive cells that co-localised with Aβ42 (Ci) and periplaque were all TMEM119-labelled cells that did not co-localise with Aβ42 (Cii).

Note that standard analysis methods for gene expression have mostly been developed for analysis of *post mortem* human tissue. This requires complex normalisation methods to overcome the problems of heterogeneity in the genetic and environmental background of the subjects, condition of the tissue, cause of death, *post mortem* interval and other variables, depending on the source of the tissue. The situation is completely different in mice that have identical genetic backgrounds (except for the genes altered for the experiment) and controlled conditions for: environment, diet, killing of the animal, treatment of the tissue and extraction methods. Consequently, much simpler methods of normalisation are sufficient, such as using housekeeping (HK) genes. As this study is concentrating on the microglial AOIs, all data were normalised to *Actg1* and *Actb.*

### Distribution of microglia around plaques in NLF mice

As has frequently been reported, microglia cluster around plaques in both mouse models and human tissue (Itagaki et al., 1989; Matarin *et al*., 2015; Medawar et al., 2019). This is also true in the NLF mice, where we have reported that an overall increase in microglial density across the hippocampus of NLF mice only reaches statistical significance at 24 months (Benitez *et al*., 2021). In order to ensure that any change in gene expression in our spatial transcriptomics analysis was adequately corrected by the normalisation procedure, we checked that expression was not substantially affected by differences in microglial density but truly reflected gene expression per microglial cell. We thus compared the distribution of microglia around individual plaques by comparing the density of IBA1 positive cells (Figure 2A and 2B) to the expression of the microglial gene *Aif1* which codes for IBA1 (Figure 2C). The expression of *Aif1* in individual microglial cells would not be expected to increase substantially in response to plaques. As expected, when density of microglia was assessed on plaques and in concentric circles increasing by 10 μm radius out from the plaque edge, there was a substantial increase in density in the proximity of plaques. However, the increase in microglial density was only detectable at the plaque or within the first 10 μm ring around the plaque. By 20 μm, the density had returned to the level far from plaques, which was also not significantly different from the density in WT mice. Despite the >30-fold increase in density of microglia at the plaque, no significant effect of region or genotype was detected in *Aif1* expression.

**Figure 2.**
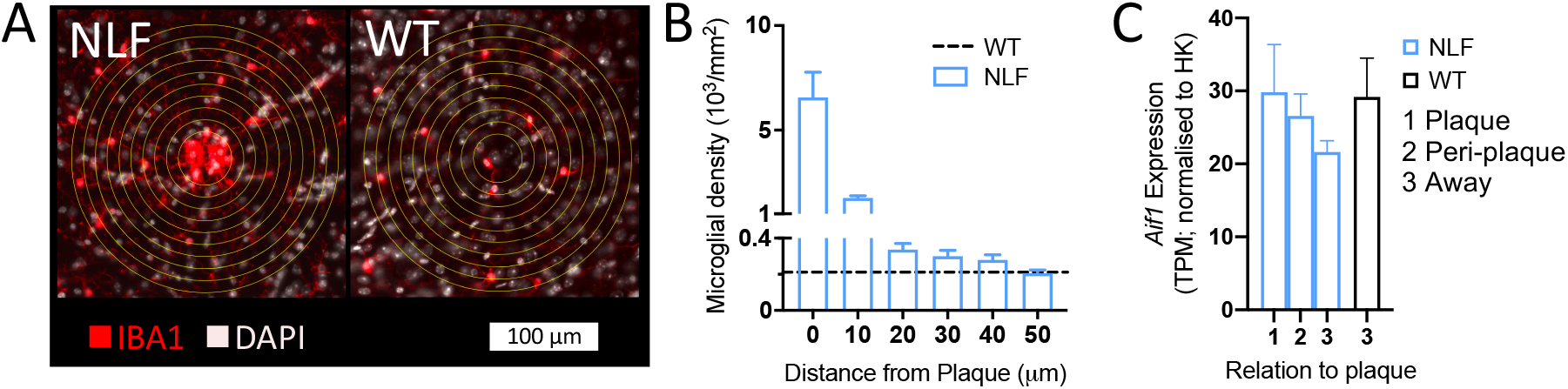
Microglial density compared to *Aif1* expression. **(A)** Microglia were labelled with an antibody against IBA1 (red), plaques with LCOs (not shown) and nuclei with DAPI (white). For NLF mice, the circumference of the plaque was identified and concentric circles were drawn at 10 μm intervals. Microglia were counted in the inner circle and each circle of increasing size. A microglial cell touching a circle was considered to lie inside that circle and density was calculated. For WT mice, a central circle with a 10 μm radius was placed pseudo-randomly to represent similar areas assessed in the NLF mice. Microglial density was counted as for NLF mice. **(B)** Quantification of microglial density in NLF mice reveals a 30-fold higher density on the plaque compared to WT mice. **(C)** the expression of *Aif1* (which encodes IBA1) in NLF mice was not significantly increased either compared to WT mice or with distance from plaque. Data expressed as mean + SEM. n=6 mice per genotype in panel B and n=6 NLF, n=4 WT mice in panel C.

It should be noted that for all immunohistochemistry experiments the plaque has been labelled with luminescent conjugated oligothiophenes (LCO). Use of LCOs will define only deposited Aβ (Hammarstrom et al., 2010; Klingstedt et al., 2011; Klingstedt et al., 2013) and so is rather more stringent in terms of defining the plaque than the Aβ42 antibody used for the transcriptomics above which may label more diffuse Aβ around the plaque.

### Most but not all previously defined plaque-induced genes, including Trem2, show significant differences between microglia on plaques and far from plaques

As a first step in comparing the regional expression of relevant genes in relation to the position of plaques, we compared the microglial enriched expression of the genes previously defined as PIGs (Chen *et al*., 2020)(Figures 3A, 3B, and 3C).

**Figure 3.**
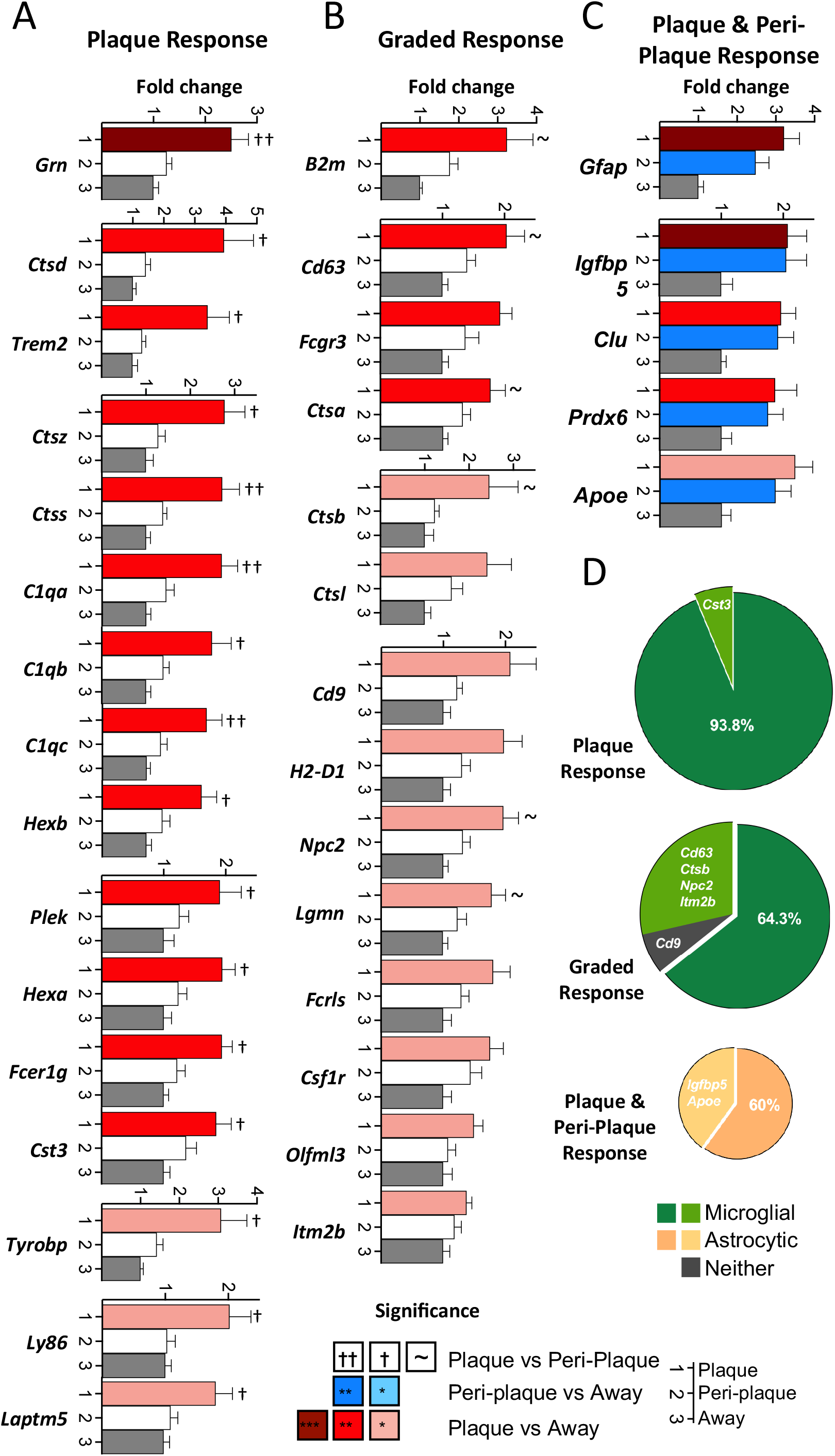
Expression of PIGs *(Overleaf)* 35 of the 55 PIGs tested in NLF mice show significant main effect of relation to plaque in a one-way ANOVA **(A)** 16 genes were only upregulated when microglia are touching plaques. **(B)** 14 genes show a graded response in which the periplaque region is not significantly different from either the plaque or away regions. **(C)** Five genes are equally upregulated in the plaque and periplaque regions, both with increased expression compared to away. For other PIGs there was no significant effect of relation to plaque.do not show. Data presented as Mean + SEM; n=6 mice. Transcripts were averaged from one to three AOI per mouse. **D**) Pie charts illustrating the percentage of genes that are identified as microglial (green), astrocytic (orange), or neither (grey). Cell specificity was initially assessed according to comparative single cell RNAseq analysis from whole brain of aged (21-22 months old) wild type mice Genes were considered specific to microglia or astrocytes if expressed > 2-fold compared to all other cells. (Ximerakis *et al*., 2019)(dark green/orange). Where no distinction was available from Ximerakis et al., the data were taken from the Barres Brain RNAseq database (Zhang *et al*., 2014) (lighter green/orange). n=6 mice. Statistical difference as indicated ***P<0.001; **/†† P<0.01; */† P<0.05, ~ indicates a strong trend (P<0.08) for plaque versus periplaque.

Of the 55 tested PIGs, expression of 35 genes showed a significant main effect of relation to plaque in the microglial AOIs (one-way Analysis of Variance). (Two pseudogenes, *Gpx4-ps & Cd63-ps,* reported as PIGs were not available in the Nanostring platform.) We divided the differentially expressed genes into 3 groups: 1. Genes only significantly raised in microglia touching plaques (Figure 3A); 2. Those that decreased more gradually, with the plaque region being significantly higher than the away region but the periplaque region not differing significantly from either (Figure 3B); and 3. Genes that were significantly upregulated in both the plaque and periplaque region compared to the away AOIs (Figure 3C). One additional PIG gene *(H2-k1)* was also of interest as it appeared to have a U-shaped response to plaque proximity being significantly increased in the periplaque area but returning to background levels of expression on the plaque (data not shown). This could suggest an effect on expression of this gene of low concentration Aβ which was lost at higher concentrations near the plaque.

Group 1 was of particular interest, as these genes are most likely responding to contact with the plaque itself, rather than to soluble Aβ or other secreted substances at a distance from the plaque. This group included *Trem2* and its downstream signalling partner *Tyrobp,* as well as several of the complement genes that have previously been implicated in AD and lysosomal genes.

Note that not all PIGs are thought to be microglial genes and so we compared the different groups in terms of previously defined relative distribution in microglia or astrocytes in single cell transcriptomic analysis comparing specific cell types across the whole brain in 21-22-month-old mice (Ximerakis *et al*., 2019). Figure 3D indicates the proportion of genes in each group that have been reported to be at least 2-fold enriched for microglia or astrocytes compared to all other cell types in whole brain. Genes that were not defined as astrocytic or microglial in the Ximerakis et al. database were further assessed using the RNAseq database (https://www.brainrnaseq.org) from the Ben Barres (Zhang et al., 2014) lab using cortical expression in young mice (Zhang *et al*., 2014). Interestingly, all genes in Group 1 and all except *Cd9* in Group 2 have been reported to be highly enriched in microglia, whereas the genes in Group 3 were reported to be enriched in astrocytes. Thus, where signal in the microglial AOIs is likely to be coming from remaining overlap of astrocytic processes, this is seen equally in the plaque and periplaque regions but occurs somewhat less in the region away from plaques. This probably reflects astrogliosis in the plaque ROIs relative to the ROIs without plaques. As shown by immunohistochemistry of GFAP labelled astrocytes, astrogliosis remains and only decreasing slightly in the periplaque regions (10-50 μm; Figure S2) unlike the on plaque specificity of microglial density (Figure 2).

### The increase in Trem2 expression in microglia touching plaques is prevented by the R47H mutation

We were particularly interested in *Trem2,* which has consistently appeared as a hub gene in our analysis and that of other groups (for example Chen *et al*., 2020; Cheng-Hathaway et al., 2018; Keren-Shaul *et al*., 2017; Matarin *et al*., 2015; Salih *et al*., 2019). In the present study, *Trem2* was amongst the PIGs which only increased in expression in microglia if they were touching plaques (Figure 3A). To investigate the effect of including the *Trem2^R47H^* risk factor mutation, the NLF mice were crossed with Trem2^R47H^ knock-in mice from Jackson Laboratories, with mice being bred to homozygosity for both the *App*^NLF^ and the *Trem2^R47H^* mutations (NLFTrem2^R47H^ mice). It has previously been reported that Trem2^R47H^ mice show a substantial decrease in *Trem2* expression in young mice (Cheng-Hathaway *et al*., 2018; Xiang et al., 2018). However, in the present study of aged mice, the difference in expression of about 20% compared to WT was not statistically significant (P=0.16). In contrast, in microglia away from plaques, although the *Trem2^R47H^* mutation did not significantly alter *Trem2* expression, the NLF mutations in *App* showed a strong trend to decreased background *Trem2* expression compared to mice with WT *App* (P=0.052; Figure 4A).

**Figure 4.**
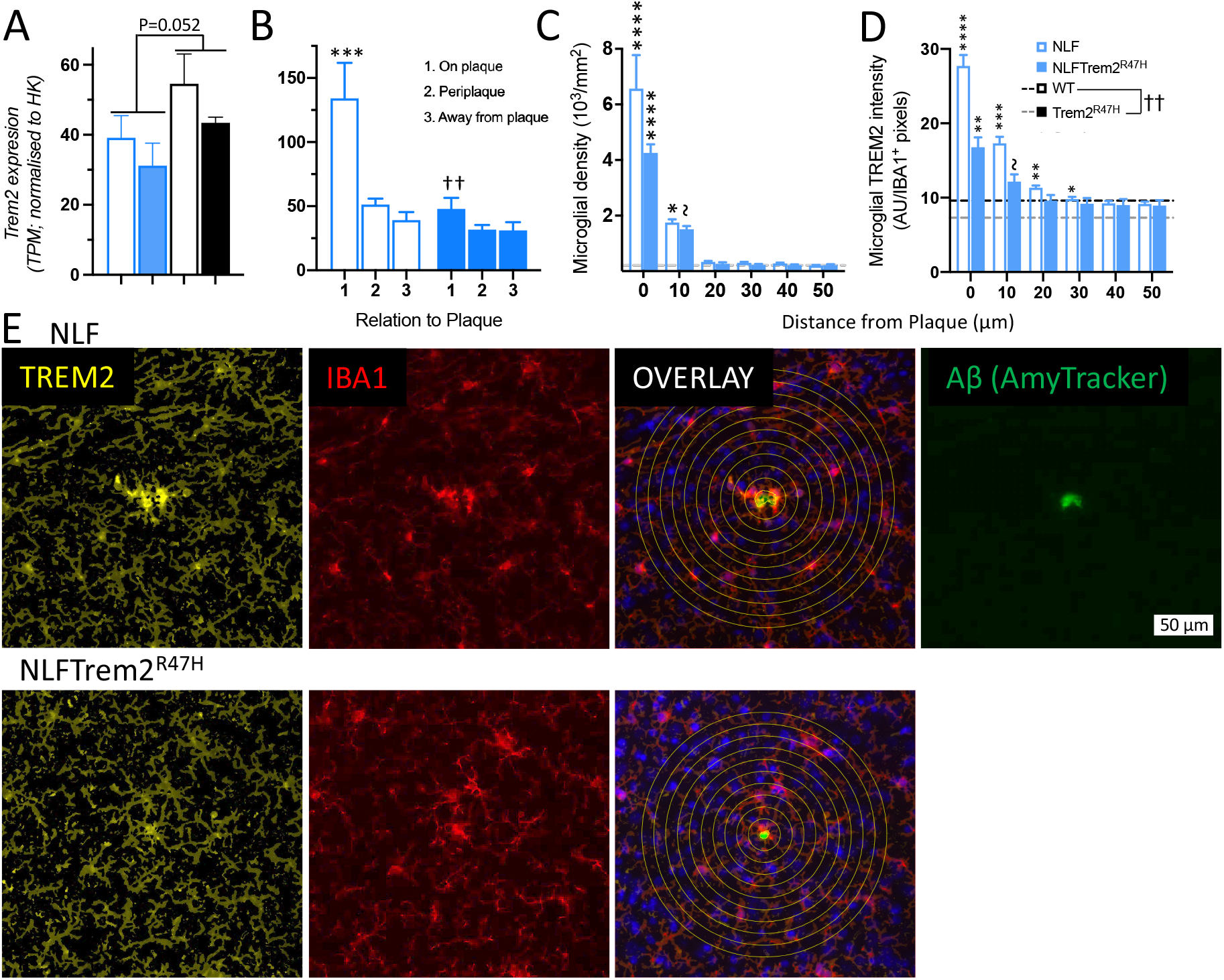
Increase microglia density at plaques is largely independent of *Trem2* genotype-dependent expression. *Trem2* expression away-from plaques. Two-way ANOVA comparing *App* genotype versus *Trem2* genotype: main effect of *App* genotype (P=0.052); no effect of *Trem2* genotype (P=0.16) and no interaction. **(B)** *Trem2* expression in NLF mice is significantly (2.6-fold) upregulated in microglial AOIs on plaques compared to periplaque AOIs. Periplaque AOIs are not significantly different from AOIs away from plaques or in WT mice. NLFTrem2^R47H^ mice show no difference on plaques compared to other regions in the same animals or compared to Trem2^R47H^ or WT mice. Two-way ANOVA between NLF and NLFTrem2^R47H^ mice: main effects of relation to plaque (P<0.01) and of genotype (P<0.01) and near significant interaction (P=0.053). *Post hoc* Sidak multiple comparison tests are shown on the graph: * within genotypes relative to NLF3 (away); † between genotype relative to NLF1 (on plaque) **(C)** Microglial density on plaques in NLFTrem2 mice is less than in NLF mice (60%, P<0.001) but is increased >20-fold compared to WT or AOIs far from plaques, despite no significant change in *Trem2* expression. Two-way ANOVA: main effect of distance from plaque (P<0.0001); interaction between distance from plaque and genotype (P=0.014). Sidak’s multiple comparison test * indicates a significant within genotype difference versus 50 μm from plaque **(D)** TREM2 protein levels reflect the distribution suggested by gene expression. Two-way ANOVA: main effects of genotype, distance from plaque and interaction (P<0.0001 in each case). Post hoc Sidak correction for multiple comparisons is shown on the graph. * compares within genotype versus 50 μm from plaque. Note that there is a significant difference in TREM2 signal of 20% between WT and Trem2^R47H^ mice (P<0.01) * within genotype versus 50 μm from plaque; n=11-12 for all genotypes, approximately equal males and females. No significant effect of sex. All data plotted as mean + SEM. ****P<0.0001; ***P<0.001; **/††P<0.01; * P<0.05 ~P<0.08 **(E)** Images of TREM2 staining comparing NLF and NLFTrem2^R47H^. TREM2 (Yellow); IBA1 microglia (red); Amytracker Aβ (green).

Comparing the effects of vicinity of plaques, as outlined above (Figure 2B), the density of microglia around the plaques is increased at the plaque in NLF mice. However, like *Trem2* expression, microglial density drops off rapidly with a significant increase remaining only in the first 10 μm ring around the plaque, where microglia may well still be in direct contact with the plaque. However, in NLFTrem2^R47H^ mice, despite there being no significant increase in *Trem2* gene expression on the plaque, the microglial density is still increased by 20-fold on plaques compared to background and similar to NLF, was back to background levels 20 μm from the plaque. Thus increased *Trem2* expression is not necessary for microglial proliferation at the plaque.

### TREM2 protein levels largely confirm plaque dependence of Trem2 expression

To assess if the remarkable fall off in *Trem2* expression away from plaques in NLF mice is reflected in a similar change in protein expression, we undertook an immunohistochemical analysis of TREM2 protein levels (Figure 4D and 4E). The analysis was restricted to TREM2 staining within IBA1+ positive pixels. Fluorescence intensity was expressed per pixel to assess the expression within microglia, independent of the density. Similar to the gene expression, there was an increase in TREM2 protein signal on the plaque in NLF mice, increasing 3-fold compared to levels away from plaques. The signal dropped rapidly to a background plateau but unlike gene expression a small significant increase was still detectable at 30 μm away from the plaque when compared to 50 μm which was not different from WT levels. Note however there is no change in microglial density in response to this more distal change in protein level.

In NLFTrem2^R47H^ mice, there was a significant increase in protein expression at the plaque and immediately adjacent to the plaque but to a significantly lower level than the NLF mice (Figure 4D and 4E). Notably, this small (1.6-fold) but significant increase in TREM2 protein did not apparently require a significant increase in gene expression in the NLFTrem2^R47H^ mice (Figure 4B), suggesting either increased translation or decreased clearance. Moreover, this related to a much greater (20-fold) increase in microglial density (Figure 4C).

### The Trem2^R47H^ risk factor mutation prevents phagocytic phenotype and induces a higher density of small plaques

The question remains as to what the microglia clustered around the plaques are doing and what is the role of TREM2 in this activity. It has repeatedly been shown that an increase in *Trem2* expression results in increased phagocytosis (Jiang et al., 2017; Liu *et al*., 2020; Takahashi et al., 2005) and that this is dependent on the *Trem2* genotype, with the R47H mutation decreasing the phagocytic activity (Kleinberger et al., 2014; Yin et al., 2016). Interestingly, impaired phagocytosis is not confirmed in macrophages from human induced pluripotent stem cells derived from patients with the *TREM2^R47H^* mutation (Cosker et al., 2021; Hall-Roberts et al., 2020). However, in agreement with previous reports (Liu *et al*., 2020), we observed the regional gene expression of *Cd68,* a marker of the phagocytic phenotype, and found that its expression mirrored that of *Trem2* and was similarly dependent on the *Trem2* genotype (Figure 5A). Hence in NLF mice, *Cd68* was only significantly upregulated in microglial-enriched AOIs that were on plaques and not in periplaque regions and this increase was completely lost in the NLFTrem2^R47H^ mice.

**Figure 5.**
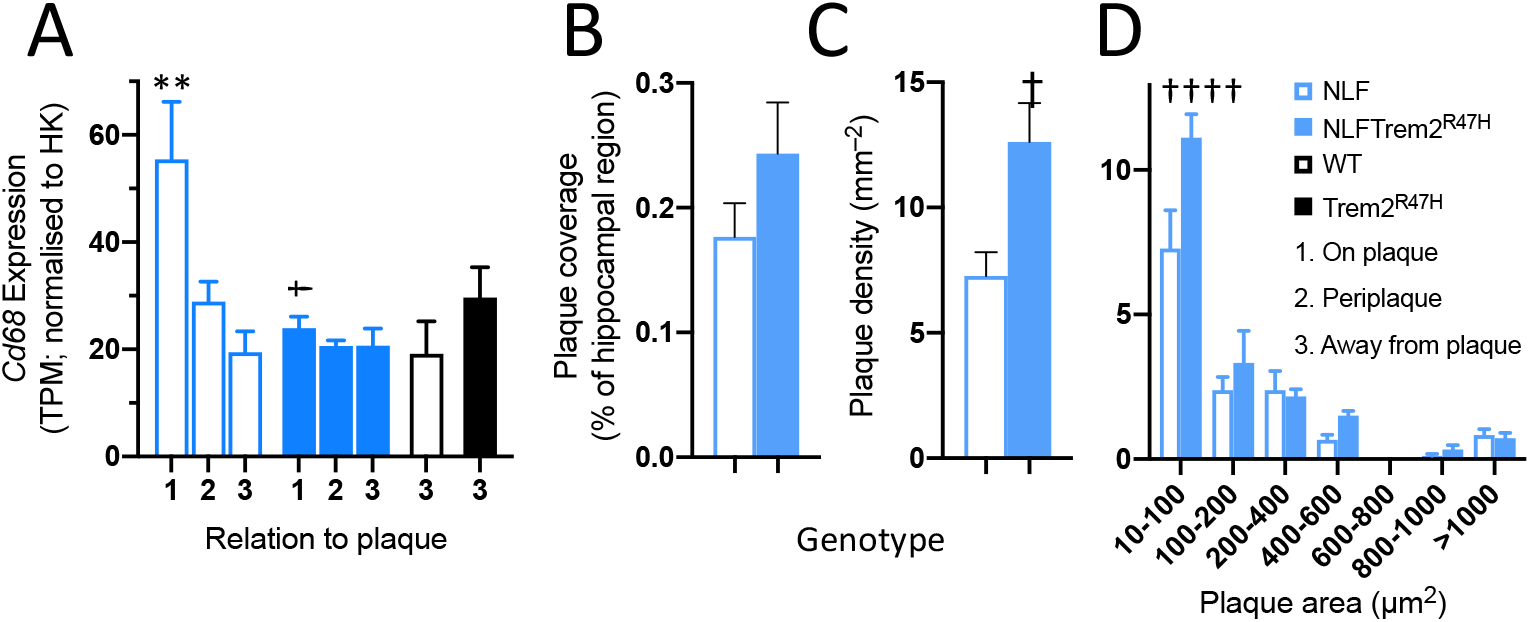
Microglial activation state and density of small plaques is affected by the *Trem2* genotype. **(A)** Expression of *Cd68* behaves similarly to *Trem2* expression, being significantly increased only in microglial AOIs touching plaques with the increased expression lost in the presence of the *Trem2^R47H^* mutation. Sidak post hoc tests are indicated by * within genotype comparison versus AOI 3 (away). † indicates across genotype comparisons with respect to same AOI, P<0.05. NLF: n=6; NLFTrem2^R47H^, WT and Trem2^R47H^, n=4 per genotype **(B-C)** While plaque coverage did not show a significant difference between NLF and NLFTrem2^R47H^ mice (B), plaque density was higher in NLFTrem2^R47H^ versus the NLF mice (C). **(D)** A plaque size-density frequency histogram revealed that the higher density of plaques was restricted to small plaques. Sidak post hoc tests across genotype † P<0.05, †††† P<0.0001. Panels B-D: n=6 mice for both genotypes. Data presented as mean + SEM in all panels. For all panels ††††P<0.0001; **/††P<0.01; † P<0.05

We then examined the plaque coverage and the number of plaques/area in the NLF and NLFTrem2^R47H^ mice. While the plaque coverage was not significantly different between the two *Trem2* genotypes (Figure 5B), the number of plaques per area was greater in the NLFTrem2^R47H^ mice (Figure 5C). We thus went on to analyse this apparent discrepancy by observing the number of plaques of different sizes. Most analysis around plaques, including the regional gene expression in the present study, concentrates on the large plaques, with cross sectional area >~100 μm^2^. These plaques dominate the plaque coverage of the brain regions studied. However, the majority of plaques, in terms of plaque number, are very small (<100 μm^2^) and this remains true throughout age or stage of plaque development in both mice and humans (Benitez *et al*., 2021; Morris et al., 1996; Murray and Dickson, 2014). When hippocampal plaque density was broken down by plaque size, the source of the difference was revealed to be entirely due to an increase of very small plaques in the NLFTrem2^R47H^ mice compared to the NLF mice (Figure 5D). This suggests that the microglia expressing high levels of *Trem2^WT^* are not effective in phagocytosing large plaques but are either instrumental in the removal of new small plaques or the microglia with the *Trem2^R47H^* mutation are causing the seeding of additional small plaques. However, in either scenario, these small plaques are not developing into additional large plaques.

### Only PIGs that depend on touching plaques for increased expression are dependent on Trem2 genotype

The regional distributions of “plaque-induced genes” (PIGs) expression in NLF mice were compared to NLFTrem2^R47H^ mice. Like *Trem2,* in six PIGs, *(C1qa, C1qc, Ctsz, Ctss, Grn, Plek),* the increased expression in microglia touching plaques was prevented by the *Trem2^R47H^* mutation (Figure 6). All of these genes fell into group 1, being only upregulated on plaques (Figure 3). This suggests that *Trem2^WT^* is instrumental in the sharp upregulation of expression of these genes on the plaque. The distribution of expression changes of the other PIGs that were increased in expression in the plaque and/or periplaque areas were not significantly affected by the *Trem2^R47H^* mutation (examples from each group in Figure S3). It is interesting to note that several of the *Trem2*-genotype-independent PIGs show expression in WT or Trem2^R47H^ mice that is close to the on plaque expression in the mice with plaques, being relatively down regulated away from plaques. The most striking example being the astrocytic gene *Clu* and to a large degree *ApoE,* both of which are GWAS risk genes (Schwartzentruber et al., 2021). This may reflect that the microglial AOI measurements of astrocytic genes are largely dependent on the degree of overlap of the remaining fine astrocytic processes (after the initial removal of the astrocyte AOIs). With astrogliosis probably showing an increase in spread of processes toward plaques rather than proliferation or migration of cell bodies (Bardehle et al., 2013; Damisah et al., 2020), the fine processes stretching towards plaques may leave fewer processes in the away from plaque areas.

**Figure 6.**
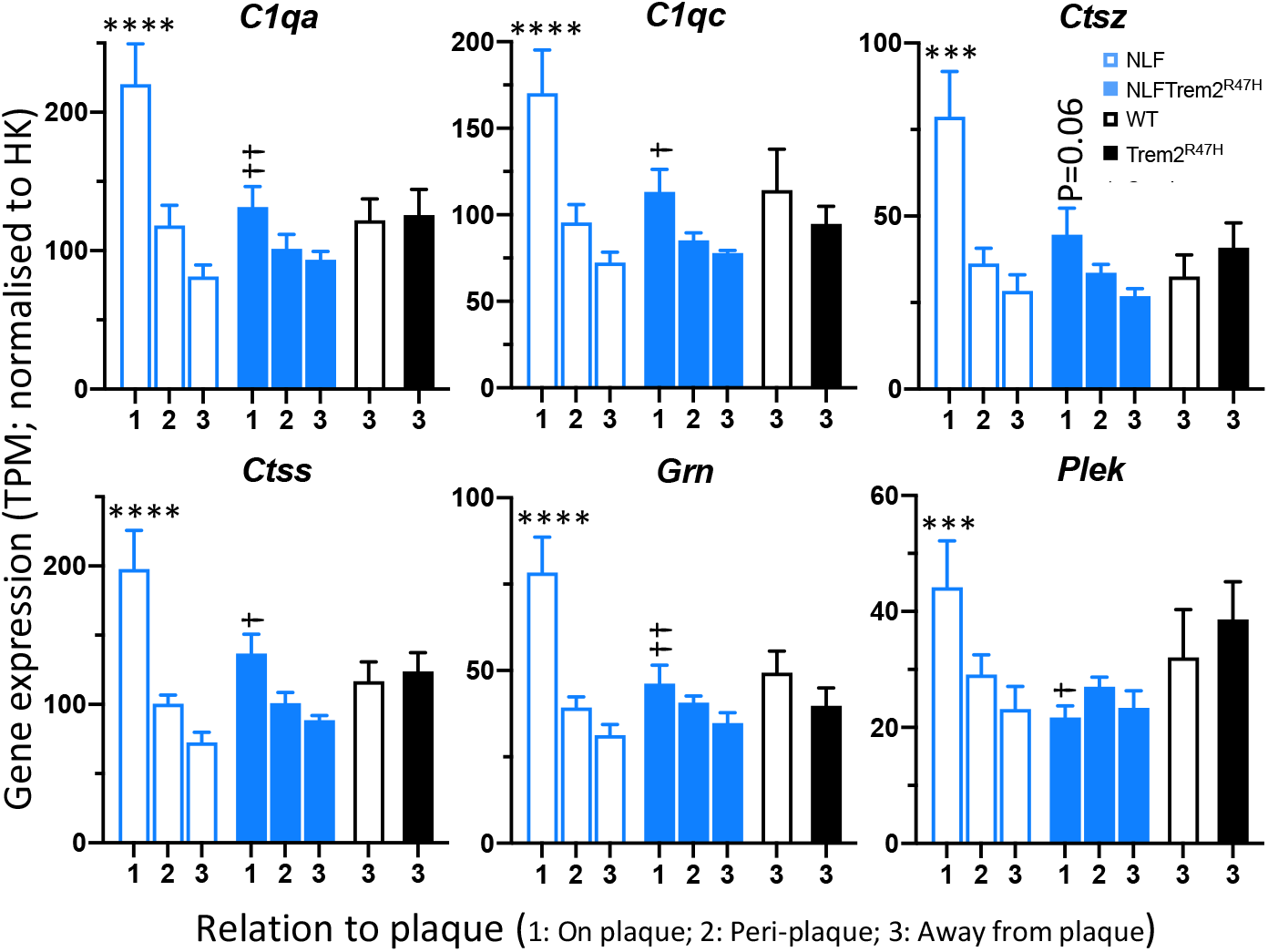
Upregulation of certain PIGs at plaques is dependent on *Trem2*^R47H^ genotype. Gene expression of six PIGs that showed a *TREM2^R47H^*-dependent upregulation in plaque AOIs only. NLF: n=6; NLFTrem2^R47H^, WT and Trem2^R47H^, n=4 per genotype. All data plotted as mean + SEM. Also see Supplementary Figure 2.

## Discussion

The observation that *Trem2* and related genes are strongly upregulated as plaques deposit in Alzheimer’s disease is well documented in studies of both mouse models (Benitez *et al*., 2021; Matarin *et al*., 2015) and human *post-mortem* tissue (Gratuze *et al*., 2018). Moreover, it has been repeatedly observed that the response of microglia around plaques is dependent on the *Trem2* genotype and is decreased with the *TREM2^R47H^* risk factor mutation (Korvatska et al., 2015) or when *Trem2* is knocked out (McQuade et al., 2020). All the previous studies in Aβ mouse models have, however, relied either on aggressive transgenic models with overexpression of APP and the other problems of transgenic technology (Sasaguri et al., 2017) or on NLGF knock-in mice in which the plaques appear by 2 months and reach an almost maximal load rapidly during early life (Benitez *et al*., 2021; Saito *et al*., 2014; Sasaguri *et al*., 2017). In the human sporadic condition, the development of plaques only starts in middle age and it is in old age that the density of plaques and other factors combine to result in the well-known clinical outcomes. Rapid deposition of plaques in the brain of a young animal may have very different effects from the gradual deposition of plaques late in life, as occurs in the human condition (Stam et al., 1986).

Although the NLF knock-in mouse carries two familial AD mutations, it represents a model of disease in which the resulting rise in Aβ leads to a gradual rise in plaque load from 9 to 24 months of age (Benitez *et al*., 2021). This results in much more subtle changes in gene expression than the related NLGF mice or the many transgenic models that preceded it and, consequently, it has been more difficult to study with standard RNAseq technology.

It is important to note that the Trem2^R47H^ knock-in mice used in this study have previously been reported to show a decrease in *Trem2* expression that was not reflected in *post mortem* tissue from people with the *TREM2^R47H^*mutation and thus was interpreted as a mouse specific artifact, possibly compromising this as a model of the human condition (Cheng-Hathaway *et al*., 2018; Liu *et al*., 2020; Xiang *et al*., 2018). However, these studies compared expression in microglia derived from very young mice or induced pluripotent stem cells to aged humans. In the aged Trem2^R47H^ knock-in mice, in the present study, the *Trem2* expression is not significantly changed compared to WT mice. It will be interesting to assess in future studies whether, rather than being mouse specific, the decreased expression may be age-related. This could suggest that effects of this mutation in increasing risk of being diagnosed in old age with AD are influenced by early changes in *Trem2* expression affecting synapses in early development.

The introduction of spatial cell-type-enriched transcriptomics has allowed more detailed and higher resolution detection of changes, within individual brain sections, at different positions in relation to the plaques than was previously possible. Under these conditions, in analysis that pairs AOIs within the same sections, we found that many, but not all, of the previously defined PIGs (Chen *et al*., 2020) were indeed increased in a plaque-related manner in NLF mice. The lack of apparent effect of vicinity of plaques in the present study of 20 of the 55 PIGs is most likely related to the different genotypes studied with Chen et al., having concentrated their study on NLGF mice that express plaques from 2 months of age and show a much stronger phenotype (Benitez *et al*., 2021; Saito *et al*., 2014). NLGF mice include an additional mutation within their Aβ sequence that results in a strong change in equilibrium of the Aβ towards deposition. Hence it may be that in older mice with heavier plaque loads that these genes would be upregulated or, alternatively that these are genes that are upregulated because of the deposition of a different Aβ structure at a much younger age. Alternatively, the difference may be related to the different methods used, with the present study relating gene expression in a cell-type enriched AOIs, dependent on their distance from plaques while the Chen study only used plaque density. Consequently, depending on the cell types in which genes are expressed, this could lead to different results between the two studies. Indeed 40% of the PIGs that did not come out in the present analysis as being differentially expressed in relation to distance from plaque had been previously identified as genes expressed in other cell types such as oligodendrocytes or epithelial cells with 3 genes *(Axl, C4a and C4b)* differentially expressed in astrocytes (Ximerakis *et al*., 2019; Zhang *et al*., 2014).

Strikingly, in the present study a subset of these genes, including *Trem2,* were only increased in the microglia AOIs that were on plaques and not in the immediately adjacent regions. Moreover, in the case of 8 of these genes, this tight regional increase in expression was not only dependent on the microglia touching the plaques but was strongly dependent on the *Trem2* genotype. This confirms the very tight relationship between these genes but refines the module to a much smaller and more clearly related group. The module includes the *C1q* complement factors that have consistently been associated with *Trem2* expression in Alzheimer’s mouse models (Matarin *et al*., 2015). However, the other genes are associated with the phagosome *(Plek)* (Brumell et al., 1999) or lysosomal function(*Ctsz, Ctss, and Grn)* (Infante et al., 2008; Lee et al., 2021a; Paushter et al., 2018; Tjondrokoesoemo et al., 2016). All of these genes are linked with the dependence on the *Trem2* genotype suggesting that *Trem2^WT^* is the essential controller. The observation that they only upregulate in microglia touching plaques, suggests that the TREM2 ligand that sets this chain in motion is hydrophobic and so does not diffuse away from the plaque. A possible candidate would be a phospholipid, such as phosphoserine that has been suggested to be an “eat me” signals on damaged synapses (Kober and Brett, 2017; Scott-Hewitt et al., 2020). This is consistent with a role for microglia in removing damaged synapses to limit the spread of damage along axons or dendrites (Edwards, 2019). The cascade then proceeds via C1Q and the complement pathway leading to phagocytosis involving pleckstrin and lysosomal removal of waste involving the cathepsins, NPC2 and progranulin. In principle this involves activation of proteins and does not need to involve gene expression if sufficient protein is already expressed by the microglia. However, the increased expression of all these genes suggests that these may be rate limiting steps in the process. This is further underlined by the fact that some increase in protein and considerable microglial proliferation occur in NLFTrem2^R47H^ knock-in mice, despite a lack of differential expression of *Trem2* at the plaque; the modest increase in translation of protein being apparently sufficient.

The link between the PIGs above leading to the phagocytic phenotype of microglia is further supported by the expression of the phagocytic marker CD68 (Zotova et al., 2011) which, although not included in the genes reported to be plaque-induced by Chen et al, shows the same behaviour being only upregulated on plaques and dependent on the *Trem2*^WT^ genotype.

An alternative hypothesis for the role of this phagocytic pathway, not mutually exclusive with the above, suggests that the role of plaque-related microglial phagocytosis relates to removal of Aβ (Yu and Ye, 2015). In this context, several studies have investigated the effect of removing microglia using inhibitors of the CSF1 receptor (Benitez *et al*., 2021; Dagher et al., 2015; Spangenberg et al., 2019; Spangenberg et al., 2016). Together, these studies suggest that microglia may have a role in seeding new plaques but that the overall plaque load, once established, is little affected by the removal of microglia (but see, for example, Zhao et al., 2017). In the present study, we found a notable effect of the *Trem2* genotype in that NLFTrem2^R47H^ mice showed a selective increase in the number of very small plaques compared to NLF mice, while larger plaques were unaffected. This is largely consistent with the findings above. It seems that the activation of microglia expressing *Trem2^WT^* may be involved in the removal of very small plaques, but once the plaques grow past about 100 μm^2^ they are no longer effective in their removal. An alternative explanation could be that the *Trem2^R47H^* mutation results in constant seeding of plaques either *de novo* (Friedrich et al., 2010) or by small plaques breaking off from the larger plaques. However, it seems that these extra plaques are not destined to grow into larger plaques. It should also be noted that if Aβ is the ligand triggering the response, that soluble Aβ would be expected to be present at a distance from the plaque in NLF mice.

## Conclusion

Use of spatial cell-enriched transcriptomics has allowed analysis of plaque-related gene expression in a mouse model that mimics the slow and age-related deposition in plaques that is seen in Alzheimer’s disease. The findings in the present study confirm the basic principles of much that has been shown in more aggressive models in which plaque load increases rapidly over early life (for example Chen *et al*., 2020; Matarin *et al*., 2015). However, it refines the effects to a smaller group of genes closely related to *Trem2* expression and dependent on the *Trem2* genotype. The comparison of the dependence of the expression of these genes on microglia touching plaques and the differential dependence on *Trem2* genotype of microglial density, strongly suggests that the increase of microglial density is, at most, marginally dependent on *Trem2* expression. Moreover, the increase in expression of this group of genes appears to depend on a hydrophobic ligand that does not diffuse far from plaques.

## Materials and Methods

### Mice

Male and female *App*^NLF/NLF^ knock-in mice (NLF), *App*^NLF/NLF^ knock-in mice harbouring the *Trem2^R47H/R47H^* mutation (NLFTrem2^R47H^) and age-matched C57BL/6J (WT) and *Trem2*^R47H/R47H^ (Trem2^R47H^) control mice were used throughout the study. No statistically significant sex differences were observed. All mice were bred in the UCL Biological Services Unit. Original breeders for the NLF mice were obtained from Riken Japan (Saito *et al*., 2014) and the Trem2^R47H^ mice from Jackson Laboratories (stock #027918). Same sex littermates were group-housed (2–5 mice) with an *ad libitum* supply of food and water, under a 12-hour light/dark photoperiod, at a controlled temperature and humidity. Experiments were performed in accordance with the UK Animal (Scientific Procedures) Act, 1986 and following local ethical review.

### Genotyping

Ear, tail or brain biopsy tissue was processed by either in-house UCL or Transnetyx (Cordova, TN, USA) genotyping services to determine the presence of the knock-in genes.

### Tissue extraction

Animals were decapitated and the brain rapidly extracted on ice and bisected in the sagittal plane. One hemisphere was drop-fixed in 10% formalin at 4°C overnight, then transferred to a 30% sucrose, 0.02% sodium azide in phosphate-buffered saline (PBS) solution for long-term storage at 4°C. The other hemisphere was either snap frozen and processed for RNA extraction or used in other experiments not presented in the current study.

### Sectioning

A portion of the dorsal cortex was removed from the formalin-fixed hemispheres, to orient the sections such that the maximal number of sections transverse to the long axis of the hippocampus could be obtained. A total of 36 serial sections containing transverse ventral-medial hippocampus were prepared at 30 μm using a frozen sledge microtome (Leica SM2010R). Sections were collected into 0.02% sodium azide in 0.1 M PBS for immunohistochemistry. A block containing the hippocampus was dissected out of the remaining un-sectioned tissue then paraffin-embedded and sectioned transverse to the long axis of the hippocampus at 8 μm using a rotary microtome (Leica RM2235) and collected directly onto SuperFrost plus slides for spatial transcriptomics.

### Nanostring GeoMx Digital Spatial Profiler

The paraffin embedded sections were processed by the Nanostring Technology Access Programme (Seattle, WA, USA). Epitope retrieval and staining was performed using the Leica Bond Autostainer. In the first step, the slides were baked at 60°C for 30 minutes. Heat induced epitope retrieval was then performed with 0.1 μg/ml proteinase K (Thermo Fisher, AM2546) in Epitope Retrieval Solution 2 (Leica, AR9640) for 20 minutes at 100°C. Following this, the slides were removed from the autostainer and hybridised with whole transcriptome RNA detection probes, each conjugated to a unique photocleavable DNA oligo tag (Nanostring Technologies), overnight at 37°C. Stringent washes in equal parts 4x saline-sodium citrate (SSC) buffer and 100% formamide were performed twice for 25 minutes at 37°C followed by a final wash in 2x SSC buffer. Non-specific binding was blocked using buffer W (proprietary blocking buffer from NanoString) followed by incubation with the appropriate antibodies (mouse anti-GFAP Alexa-Fluor 488 conjugate, (Invitrogen, Cat # 53-9892-82), mouse anti-Aβ40/42 Alexa Fluor 594 conjugate (Nanostring, Cat # 121301306) and rabbit anti-TMEM119 (SynapticSystems, Cat # 400 002) alongside goat anti-rabbit Alexa Fluor 647 (Thermo Fisher, Cat # A11037)). Nuclei were counterstained (SYTO 83, Invitrogen, S11364) in buffer W for 1 hour at room temperature. Finally, the slides were washed in 2x SSC buffer.

### Region of Interest (ROI) Selection

ROIs were defined based on the morphological stains. For each NLF and NLFTrem2^R47H^ mouse two types of ROI were defined: 1. in the region of heaviest plaque load in which all astrocytes and microglia would fall within about 30 μm of a plaque and 2. (for all genotypes), a region which did not contain plaques and with borders >50 μm from any visible plaque. One section from each mouse was used to select ROIs, with 1-3 of each type of ROI per section.

### Area of Interest (AOI) selection

<u>Within the plaque containing ROI</u>, AOIs were defined as follows:

- On plaque microglia, co-localisation of TMEM119 & Aβ42;
- Periplaque microglia, TMEM119 staining not colocalised with Aβ42.
- Plaque/periplaque astrocytes, all GFAP staining.

<u>Within the ROIs without plaques</u>, AOIs were defined as follows:

- Away microglia, all TMEM119 staining
- Away astrocytes, all GFAP staining

### Library preparation and readout

RNA probe-associated DNA oligo tags were released by illuminating the chosen AOIs under ultraviolet laser exposure and aspirated into a microtiter plate, whereby each well contains the DNA oligo tags of a single AOI. The aspirates were dried and diluted in nuclease free water. DNA oligo tags were uracil removed and PCR cycled with GeoMx seq code PCR master mix (Nanostring Technologies) and GeoMx SeqCode primers (Nanostring Technologies). PCR product was AMPure bead purified initially with a 1.2x bead:sample ratio followed by a 1.2x bead:resuspended sample ratio. Samples were then sequenced on an Illumina NextSeq 2000. Filtering and demultiplexing were carried out according to (Khan et al., 2021).

### Analysis

Deduplicated counts were normalised to the average *Actb* and *Actg1* count as follows. For each AOI, the count for each gene was divided by the mean of the *Actb* and *Actg1* counts in that AOI and the normalised value multiplied by the mean count for *Actb* and *Actg1* across all AOIs, resulting in a normalised expression value that was comparable to the level of the raw data. Where this fell below 15 the ROI was discarded. Where >1 AOI was collected from a section for the same category (plaque, periplaque or away) the data were averaged giving n=1 per section to avoid pseudo-replication.

### Histology

Free-floating 30 μm sections were washed once in PBS for 10 minutes. The sections then underwent antigen retrieval in SSC buffer (10mM, pH 9.0) for 30 minutes at 80°C. The tissues were permeabilised by washing three times for 10 minutes in 0.3% Triton X-100 in PBS (PBST). Non-specific binding was blocked for 1 hour in 10% donkey serum, followed by incubation with the appropriate primary antibodies in blocking solution at 4°C overnight (1:1000 rabbit anti-IBA1 (FujiFilm Wako Chemicals, Cat # 019-19741) & 1:500 sheep anti-TREM2 (R&D Systems, Cat # AF1729)). The next day sections were washed three times for 10 minutes in PBST followed by a 2-hour incubation in the dark with secondary antibodies in blocking solution (1:1000 donkey anti-rabbit Alexa Fluor 594 (ThermoFisher Scientific, Cat # A32 and 1:1000 donkey anti-sheep Alexa Fluor 647 (Abcam, Cat # AB150179)). Finally, nuclei were counterstained with 4’,6-diamidino-2-phenylindole (DAPI) and plaques labelled with Amytracker520 (Ebba Biotech, Sweden).

### Imaging for plaque counts and immunohistochemistry

Epiflourescent photomicrographs of whole cross-sectional hippocampal regions were serial scanned under a 20x objective at constant light, gain and exposure settings using an EVOS® Fl Auto Cell imaging microscope. Three sections per animal were imaged. For all analyses, data from replicates from a single animal were averaged for n=1 estimate/animal.

### Histological Analysis

#### Protein intensity

Analysis was carried out using custom written semi-automated macros within ImageJ. The macros are available at: *https://doi.org/10.5281/zenodo.5847431*

##### 1. Plaque thresholding macro

The hippocampal region within an image was selected by manual tracing and subsequently converted to 8-bit colour and individually thresholded to account for background stain variability. The threshold for each section was established as follows:

As the Image J threshold setting is increased, the area measured decreases. By plotting a range of 40 increasing threshold settings against the associated thresholded area, as a percentage of total area, the points form an exponential decay to a 0% asymptote. When the final 40% of the values are fitted with a straight line, the intersection of this line with the data points is used for the threshold for that section. Particles <10 μm^2^ were considered as noise and excluded from the analysis.

##### 2. Plaque concentric circles macro

For each plaque within the selected hippocampal region, concentric circles were drawn outwards with increasing 10 μm radii from the plaque, reaching a final circle with a radius of the average plaque radius plus 100 μm. Where two plaques were distanced <200 μm apart, concentric circles terminated at the nearest 10 μm to the halfway point to avoid overlapping data. As most plaques in the plaque ROIs were not more than 100 μm apart data are shown up to 50 μm from the plaque edge.

##### 3. Concentric circles macro for controls

For images from WT and NLFTrem2^R47H^ mice, ten randomly placed circles each with a 20 μm radius, were positioned within the hippocampus and to imitate the measurement in relation to plaques above, circles were drawn outwards with incrementing 10 μm radii reaching distance of 100 μm from the inner circle.

##### 4. Protein intensity macro

Fluorescence intensity within the regions identified as microglia (AU/pixels) was calculated for plaque regions and radiating concentric rings.

### Microglial Density

Following running of the macros, microglial number (cells coexpressing IBA1 and DAPI) was manually counted for each concentric ring using Adobe Photoshop 2020 and density calculated by dividing by ring area.

### Statistical Analysis

All statistical analyses were performed using GraphPad Prism 9. Two-way ANOVAs were followed by Tukey-corrected post hoc analyses when a significant interaction was obtained. All sample sizes represent numbers of animals; the mean of technical repeats on samples from the same animal were calculated prior to analysis.

## Funding

This work was largely funded by a grant from Alzheimer’s Research UK (FAE and DMC) and The Cure Alzheimer’s Fund (FAE and JH). JH is supported by the Dolby Foundation. His work was partly funded by the UK DRI, which receives its funding from the DRI Ltd, funded by the UK Medical Research Council. JW is funded by the Swedish Alzheimer’s Foundation, KSV was funded by a studentship from the UK Medical Research Council.

## Author Contributions

Conceptualisation: FAE, JW

Methodology: FAE, DMC, JW, AV, RJ

Investigation: JW, EW, RJ, KSV, SD, AB, S-LJP, FL

Visualization: FAE, JW, DMC

Supervision: DMC, FAE, TT

Writing-original draft: FAE, JW

Writing—review and editing: JW, JHardy, DMC, FAE (all authors)

Funding – FAE, DMC, JHardy, JHanrieder

## Competing interests

AV and S-LJP were employed by Nanostring Technologies.

All other authors declare they have no competing interests.

## Data and materials availability

All data available on request to corresponding author. The full transcriptomics database is under preparation for publication. The mice are under MTAs from Riken *(APP^NLF/NLF^)* and Jackson Laboratories (*Trem2^R47H/R47H^*).

## References

Bardehle, S., Kruger, M., Buggenthin, F., Schwausch, J., Ninkovic, J., Clevers, H., Snippert, H.J., Theis, F.J., Meyer-Luehmann, M., Bechmann, I., et al. (2013). Live imaging of astrocyte responses to acute injury reveals selective juxtavascular proliferation. Nat Neurosci 16, 580–586.

Benitez, D.P., Jiang, S., Wood, J., Wang, R., Hall, C.M., Peerboom, C., Wong, N., Stringer, K.M., Vitanova, K.S., Smith, V.C., et al. (2021). Knock-in models related to Alzheimer’s disease: synaptic transmission, plaques and the role of microglia. Mol Neurodegener 16, 47.

Brumell, J.H., Howard, J.C., Craig, K., Grinstein, S., Schreiber, A.D., and Tyers, M. (1999). Expression of the protein kinase C substrate pleckstrin in macrophages: association with phagosomal membranes. J Immunol 163, 3388–3395.

Chen, W.T., Lu, A., Craessaerts, K., Pavie, B., Sala Frigerio, C., Corthout, N., Qian, X., Lalakova, J., Kuhnemund, M., Voytyuk, I., et al. (2020). Spatial Transcriptomics and In Situ Sequencing to Study Alzheimer’s Disease. Cell 182, 976–991 e919.

Cheng-Hathaway, P.J., Reed-Geaghan, E.G., Jay, T.R., Casali, B.T., Bemiller, S.M., Puntambekar, S.S., von Saucken, V.E., Williams, R.Y., Karlo, J.C., Moutinho, M., et al. (2018). The Trem2 R47H variant confers loss-of-function-like phenotypes in Alzheimer’s disease. Mol Neurodegener 13, 29.

Cosker, K., Mallach, A., Limaye, J., Piers, T.M., Staddon, J., Neame, S.J., Hardy, J., and Pocock, J.M. (2021). Microglial signalling pathway deficits associated with the patient derived R47H TREM2 variants linked to AD indicate inability to activate inflammasome. Sci Rep 11, 13316.

Dagher, N.N., Najafi, A.R., Kayala, K.M., Elmore, M.R., White, T.E., Medeiros, R., West, B.L., and Green, K.N. (2015). Colony-stimulating factor 1 receptor inhibition prevents microglial plaque association and improves cognition in 3xTg-AD mice. J Neuroinflamm 12, 139.

Damisah, E.C., Hill, R.A., Rai, A., Chen, F., Rothlin, C.V., Ghosh, S., and Grutzendler, J. (2020). Astrocytes and microglia play orchestrated roles and respect phagocytic territories during neuronal corpse removal in vivo. Sci Adv 6, eaba3239.

Deczkowska, A., Keren-Shaul, H., Weiner, A., Colonna, M., Schwartz, M., and Amit, I. (2018). Disease-Associated Microglia: A Universal Immune Sensor of Neurodegeneration. Cell 173, 1073–1081.

Edwards, F.A. (2019). A Unifying Hypothesis for Alzheimer’s Disease: From Plaques to Neurodegeneration. Trends Neurosci 42, 310–322.

Friedrich, R.P., Tepper, K., Ronicke, R., Soom, M., Westermann, M., Reymann, K., Kaether, C., and Fandrich, M. (2010). Mechanism of amyloid plaque formation suggests an intracellular basis of Abeta pathogenicity. P Natl Acad Sci USA 107, 1942–1947.

Gratuze, M., Leyns, C.E.G., and Holtzman, D.M. (2018). New insights into the role of TREM2 in Alzheimer’s disease. Mol Neurodegener 13, 66.

Guerreiro, R., Wojtas, A., Bras, J., Carrasquillo, M., Rogaeva, E., Majounie, E., Cruchaga, C., Sassi, C., Kauwe, J.S., Younkin, S., et al. (2013). TREM2 variants in Alzheimer’s disease. N Engl J Med 368, 117–127.

Hall-Roberts, H., Agarwal, D., Obst, J., Smith, T.B., Monzón-Sandoval, J., Di Daniel, E., Webber, C., James, W.S., Mead, E., Davis, J.B., and Cowley, S.A. (2020). TREM2 Alzheimer’s variant R47H causes similar transcriptional dysregulation to knockout, yet only subtle functional phenotypes in human iPSC-derived macrophages. Alzheimers Res Ther 12, 151.

Hammarstrom, P., Simon, R., Nystrom, S., Konradsson, P., Aslund, A., and Nilsson, K.P. (2010). A fluorescent pentameric thiophene derivative detects in vitro-formed prefibrillar protein aggregates. Biochemistry 49, 6838–6845.

Infante, R.E., Wang, M.L., Radhakrishnan, A., Kwon, H.J., Brown, M.S., and Goldstein, J.L. (2008). NPC2 facilitates bidirectional transfer of cholesterol between NPC1 and lipid bilayers, a step in cholesterol egress from lysosomes. Proceedings of the National Academy of Sciences 105, 15287–15292.

Itagaki, S., McGeer, P.L., Akiyama, H., Zhu, S., and Selkoe, D. (1989). Relationship of microglia and astrocytes to amyloid deposits of Alzheimer disease. Journal of neuroimmunology 24, 173–182.

Jiang, T., Wan, Y., Zhang, Y.D., Zhou, J.S., Gao, Q., Zhu, X.C., Shi, J.Q., Lu, H., Tan, L., and Yu, J.T. (2017). TREM2 Overexpression has No Improvement on Neuropathology and Cognitive Impairment in Aging APPswe/PS1dE9 Mice. Mol Neurobiol 54, 855–865.

Jonsson, T., Stefansson, H., Steinberg, S., Jonsdottir, I., Jonsson, P.V., Snaedal, J., Bjornsson, S., Huttenlocher, J., Levey, A.I., Lah, J.J., et al. (2013). Variant of TREM2 associated with the risk of Alzheimer’s disease. N Engl J Med 368, 107–116.

Keren-Shaul, H., Spinrad, A., Weiner, A., Matcovitch-Natan, O., Dvir-Szternfeld, R., Ulland, T.K., David, E., Baruch, K., Lara-Astaiso, D., Toth, B., et al. (2017). A Unique Microglia Type Associated with Restricting Development of Alzheimer’s Disease. Cell 169, 1276–1290 e1217.

Kleinberger, G., Yamanishi, Y., Suarez-Calvet, M., Czirr, E., Lohmann, E., Cuyvers, E., Struyfs, H., Pettkus, N., Wenninger-Weinzierl, A., Mazaheri, F., et al. (2014). TREM2 mutations implicated in neurodegeneration impair cell surface transport and phagocytosis. Sci Transl Med 6, 243ra286.

Klingstedt, T., Aslund, A., Simon, R.A., Johansson, L.B., Mason, J.J., Nystrom, S., Hammarstrom, P., and Nilsson, K.P. (2011). Synthesis of a library of oligothiophenes and their utilization as fluorescent ligands for spectral assignment of protein aggregates. Org Biomol Chem 9, 8356–8370.

Klingstedt, T., Blechschmidt, C., Nogalska, A., Prokop, S., Haggqvist, B., Danielsson, O., Engel, W.K., Askanas, V., Heppner, F.L., and Nilsson, K.P. (2013). Luminescent conjugated oligothiophenes for sensitive fluorescent assignment of protein inclusion bodies. Chembiochem 14, 607–616.

Kober, D.L., and Brett, T.J. (2017). TREM2-Ligand Interactions in Health and Disease. J Mol Biol 429, 1607–1629.

Korvatska, O., Leverenz, J.B., Jayadev, S., McMillan, P., Kurtz, I., Guo, X., Rumbaugh, M., Matsushita, M., Girirajan, S., Dorschner, M.O., et al. (2015). R47H Variant of TREM2 Associated With Alzheimer Disease in a Large Late-Onset Family: Clinical, Genetic, and Neuropathological Study. JAMA Neurol 72, 920–927.

Kulkarni, B., Kumar, D., Cruz-Martins, N., and Sellamuthu, S. (2021). Role of TREM2 in Alzheimer’s Disease: A Long Road Ahead. Mol Neurobiol 58, 5239–5252.

Lee, J.Y., Marian, O.C., and Don, A.S. (2021a). Defective Lysosomal Lipid Catabolism as a Common Pathogenic Mechanism for Dementia. Neuromolecular Med 23, 1–24.

Lee, S.H., Meilandt, W.J., Xie, L., Gandham, V.D., Ngu, H., Barck, K.H., Rezzonico, M.G., Imperio, J., Lalehzadeh, G., Huntley, M.A., et al. (2021b). Trem2 restrains the enhancement of tau accumulation and neurodegeneration by beta-amyloid pathology. Neuron 109, 1283–1301 e1286.

Liu, W., Taso, O., Wang, R., Bayram, S., Graham, A.C., Garcia-Reitboeck, P., Mallach, A., Andrews, W.D., Piers, T.M., Botia, J.A., et al. (2020). Trem2 promotes anti-inflammatory responses in microglia and is suppressed under pro-inflammatory conditions. Hum Mol Genet 29, 3224–3248.

Matarin, M., Salih, D.A., Yasvoina, M., Cummings, D.M., Guelfi, S., Liu, W., Nahaboo Solim, M.A., Moens, T.G., Paublete, R.M., Ali, S.S., et al. (2015). A genome-wide gene-expression analysis and database in transgenic mice during development of amyloid or tau pathology. Cell Rep 10, 633–644.

McQuade, A., Kang, Y.J., Hasselmann, J., Jairaman, A., Sotelo, A., Coburn, M., Shabestari, S.K., Chadarevian, J.P., Fote, G., Tu, C.H., et al. (2020). Gene expression and functional deficits underlie TREM2-knockout microglia responses in human models of Alzheimer’s disease. Nat Commun 11, 5370.

Medawar, E., Benway, T.A., Liu, W., Hanan, T.A., Haslehurst, P., James, O.T., Yap, K., Muessig, L., Moroni, F., Nahaboo Solim, M.A., et al. (2019). Effects of rising amyloidbeta levels on hippocampal synaptic transmission, microglial response and cognition in APPSwe/PSEN1M146V transgenic mice. EBioMedicine 39, 422–435.

Morris, J.C., Storandt, M., McKeel, D.W., Jr., Rubin, E.H., Price, J.L., Grant, E.A., and Berg, L. (1996). Cerebral amyloid deposition and diffuse plaques in “normal” aging: Evidence for presymptomatic and very mild Alzheimer’s disease. Neurology 46, 707–719.

Murray, M.E., and Dickson, D.W. (2014). Is pathological aging a successful resistance against amyloid-beta or preclinical Alzheimer’s disease? Alzheimers Res Ther 6, 24.

Paushter, D.H., Du, H., Feng, T., and Hu, F. (2018). The lysosomal function of progranulin, a guardian against neurodegeneration. Acta Neuropathol 136, 1–17.

Saito, T., Matsuba, Y., Mihira, N., Takano, J., Nilsson, P., Itohara, S., Iwata, N., and Saido, T.C. (2014). Single App knock-in mouse models of Alzheimer’s disease. Nat Neurosci 17, 661–663.

Salih, D.A., Bayram, S., Guelfi, S., Reynolds, R.H., Shoai, M., Ryten, M., Brenton, J.W., Zhang, D., Matarin, M., Botia, J.A., et al. (2019). Genetic variability in response to amyloid beta deposition influences Alzheimer’s disease risk. Brain Commun 1, fcz022.

Sasaguri, H., Nilsson, P., Hashimoto, S., Nagata, K., Saito, T., De Strooper, B., Hardy, J., Vassar, R., Winblad, B., and Saido, T.C. (2017). APP mouse models for Alzheimer’s disease preclinical studies. The EMBO journal 36, 2473–2487.

Schwartzentruber, J., Cooper, S., Liu, J.Z., Barrio-Hernandez, I., Bello, E., Kumasaka, N., Young, A.M.H., Franklin, R.J.M., Johnson, T., Estrada, K., et al. (2021). Genome-wide meta-analysis, fine-mapping and integrative prioritization implicate new Alzheimer’s disease risk genes. Nature genetics 53, 392–402.

Scott-Hewitt, N., Perrucci, F., Morini, R., Erreni, M., Mahoney, M., Witkowska, A., Carey, A., Faggiani, E., Schuetz, L.T., Mason, S., et al. (2020). Local externalization of phosphatidylserine mediates developmental synaptic pruning by microglia. The EMBO journal 39, e105380.

Spangenberg, E., Severson, P.L., Hohsfield, L.A., Crapser, J., Zhang, J., Burton, E.A., Zhang, Y., Spevak, W., Lin, J., Phan, N.Y., et al. (2019). Sustained microglial depletion with CSF1R inhibitor impairs parenchymal plaque development in an Alzheimer’s disease model. Nat Commun 10, 3758.

Spangenberg, E.E., Lee, R.J., Najafi, A.R., Rice, R.A., Elmore, M.R., Blurton-Jones, M., West, B.L., and Green, K. N. (2016). Eliminating microglia in Alzheimer’s mice prevents neuronal loss without modulating amyloid-β pathology. Brain 139, 1265–1281.

Stam, F.C., Wigboldus, J.M., and Smeulders, A.W. (1986). Age incidence of senile brain amyloidosis. Pathol Res Pract 181, 558–562.

Takahashi, K., Rochford, C.D., and Neumann, H. (2005). Clearance of apoptotic neurons without inflammation by microglial triggering receptor expressed on myeloid cells-2. J Exp Med 201, 647–657.

Tjondrokoesoemo, A., Schips, T.G., Sargent, M.A., Vanhoutte, D., Kanisicak, O., Prasad, V., Lin, S.C., Maillet, M., and Molkentin, J.D. (2016). Cathepsin S Contributes to the Pathogenesis of Muscular Dystrophy in Mice. The Journal of biological chemistry 291, 9920–9928.

Xiang, X., Piers, T.M., Wefers, B., Zhu, K., Mallach, A., Brunner, B., Kleinberger, G., Song, W., Colonna, M., Herms, J., et al. (2018). The Trem2 R47H Alzheimer’s risk variant impairs splicing and reduces Trem2 mRNA and protein in mice but not in humans. Mol Neurodegener 13, 49.

Ximerakis, M., Lipnick, S.L., Innes, B.T., Simmons, S.K., Adiconis, X., Dionne, D., Mayweather, B.A., Nguyen, L., Niziolek, Z., Ozek, C., et al. (2019). Single-cell transcriptomic profiling of the aging mouse brain. Nat Neurosci 22, 1696–1708.

Yin, J., Liu, X., He, Q., Zhou, L., Yuan, Z., and Zhao, S. (2016). Vps35-dependent recycling of Trem2 regulates microglial function. Traffic 17, 1286–1296.

Yu, Y., and Ye, R.D. (2015). Microglial Abeta receptors in Alzheimer’s disease. Cell Mol Neurobiol 35, 71–83.

Zhang, Y., Chen, K., Sloan, S.A., Bennett, M.L., Scholze, A.R., O’Keeffe, S., Phatnani, H.P., Guarnieri, P., Caneda, C., Ruderisch, N., et al. (2014). An RNA-sequencing transcriptome and splicing database of glia, neurons, and vascular cells of the cerebral cortex. J Neurosci 34, 11929–11947.

Zhao, R., Hu, W., Tsai, J., Li, W., and Gan, W.B. (2017). Microglia limit the expansion of beta-amyloid plaques in a mouse model of Alzheimer’s disease. Mol Neurodegener 12, 47.

Zotova, E., Holmes, C., Johnston, D., Neal, J.W., Nicoll, J.A., and Boche, D. (2011). Microglial alterations in human Alzheimer’s disease following Abeta42 immunization. Neuropathol Appl Neurobiol 37, 513–524.

