## Supplemental Figures for "Upregulation of *Trem2* expression occurs exclusively on microglial contact with plaques"

**Supplementary Figures**

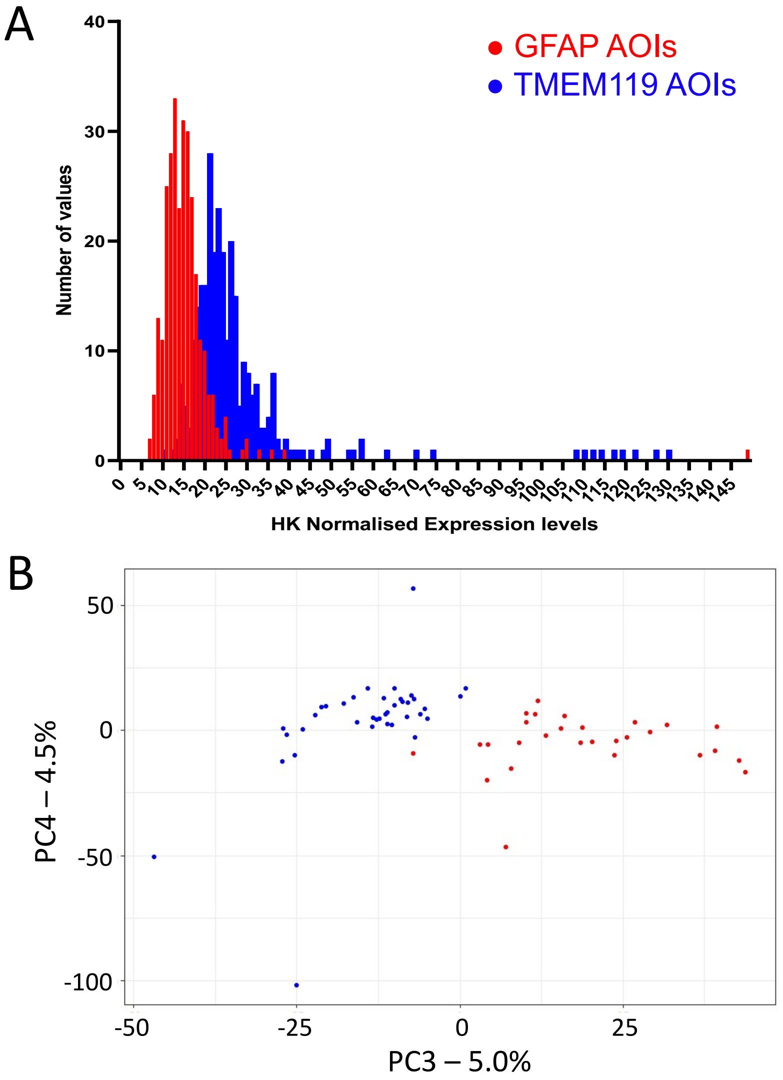


**Supplementary Figure 1. Separation of genes expressed in astrocytic and microglial AOIs.**

**(A)** Spatial transcriptomics expression within microglial (blue) and astrocytic (red) AOIs of WT mice for genes reported to be selectively expressed in microglia by at least 8-fold compared to all other genes in bulk tissue (Ximerakis *et al.*, 2019)**.** While there is some overlap considerable separation is evident. n=4 **(B)** Principal components analysis showing separation of expression of HK normalised genes in microglial (TMEM119, blue) and astrocytic (GFAP, red) enriched AOIs. n=66 AOIs from n=18 mice.

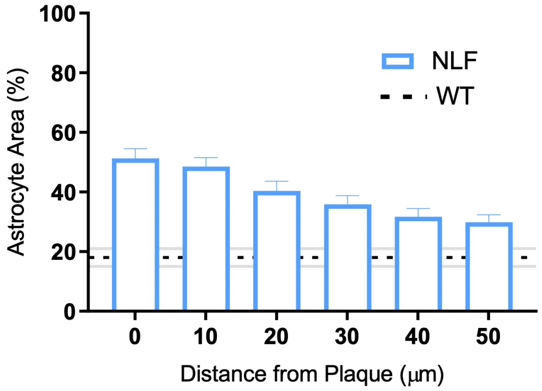


**Supplementary Figure 2. Immunohistochemical analysis of GFAP labelled astrocytes showing percentage coverage of areas in concentric rings at increasing distances from plaques.**

Unlike microglia, astrocytes have processes spreading a considerable distance. Similar methods were used to counting microglia, with concentric rings at increasing distances from plaques but rather than counting astrocytes, the area covered by astrocytic processes was compared. A two-way ANOVA showed a main effect of distance from plaque (P<0.0001) and of genotype (P<0.001) and an interaction between genotype and relation to plaque (P<0001) reflecting the gradual decline in astrocyte area at increasing distances from plaque with no change in WT. Hence astrocytic processes increase in area in response to plaques but only change gradually remaining significantly different from WT at 50 μm from the plaque. n=6 mice. Data represent mean + SEM. For WT mean across all concentric circles (black line) SEM (grey lines).

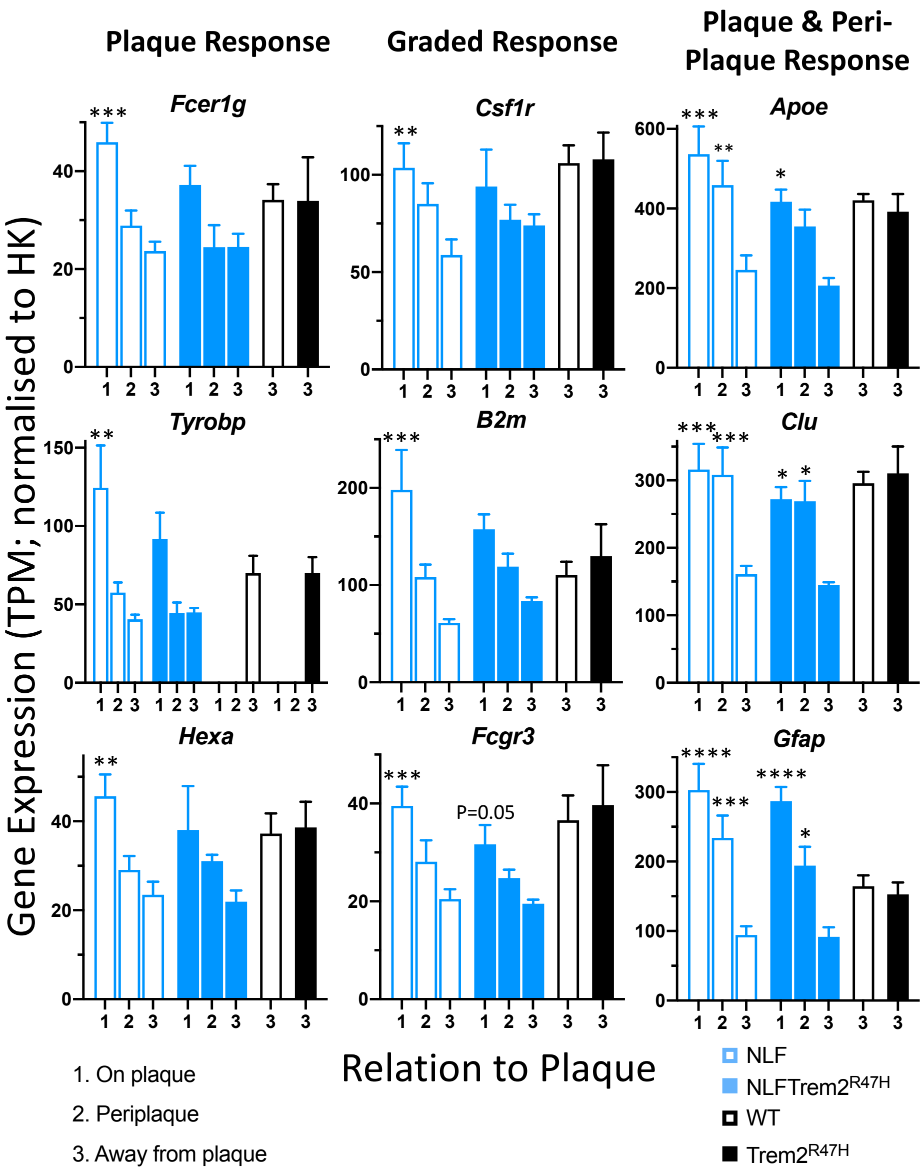


**Supplementary Figure 3. PIGs independent of *Trem2* genotype.**

PIGs that showed little dependence on *Trem2* still fell into three groups based on their expression patters in relation to plaque. *Left*: Genes that were upregulated in both NLF and NLFTrem2$^{R47H}$ mice only at the plaque. *Centre:* Genes that showed a graded response with distance from plaque. *Right:* Genes that were upregulated both on the plaque and periplaque AOIs. Note the expression levels in the WT and Trem2$^{R47H}$ mice, which, in some cases, are closer to plaque or periplaque levels than to the away levels. NLF: n=6; NLFTrem2$^{R47H}$, WT and Trem2$^{R47H}$, n=4 per genotype.
